# The deubiquitinating enzyme Otu1 releases substrates from the conserved initiation complex of the Cdc48/p97 ATPase for proteasomal degradation

**DOI:** 10.1101/2025.11.08.687396

**Authors:** Hao Li, Haipeng Guan, Tom A. Rapoport

## Abstract

Many eukaryotic proteins are modified with a polyubiquitin chain and then recruited to either the Cdc48 ATPase (p97 or VCP in mammals) or the 26S proteasome by conserved cofactors. They can then shuttle between the Cdc48 ATPase and the 26S proteasome before being degraded. How substrates avoid being trapped on the Cdc48 ATPase complex is incompletely understood, as they can undergo repeated cycles of translocation through the ATPase pore. Here, we show that the deubiquitinating enzyme (DUB) Otu1 (Yod1 in mammals) can break this futile cycle. Otu1 trims the ubiquitin chain of the substrate before its translocation through the Cdc48 pore is initiated, allowing transfer to the proteasome and subsequent degradation. A cryo-EM structure shows that the mammalian homolog Yod1 binds to p97 simultaneously with other Cdc48 cofactors. As in the yeast system, polypeptide translocation through the ATPase pore is initiated by the unfolding of a ubiquitin molecule, suggesting that the mechanism of substrate processing is conserved in all eukaryotes.

## Introduction

Regulated protein degradation in eukaryotes generally begins with the attachment of a polyubiquitin chain^1^. If the target protein contains a flexible segment, it can directly be degraded by the 26S proteasome; the segment inserts into the pore of the 19S regulatory particle and initiates substrate transfer into the proteolytic chamber of the 20S core particle^2^. In contrast, well-folded proteins require initial unfolding, a step mediated by the ATPase Cdc48 in yeast and its homolog p97 in mammals (also called VCP)^3^. Cdc48/p97 consists of an N-terminal (N) domain and two ATPase domains, D1 and D2. Six Cdc48/p97 molecules assemble into a double-ring hexameric structure with a central pore. Substrate is recruited to one side of the Cdc48/p97 double-ring (the cis side) through binding of the attached ubiquitin chain to a cofactor, the heterodimeric Ufd1-Npl4 (UN) complex. A cryogenic electron microscopy (cryo-EM) structure of the yeast Cdc48-UN complex with polyubiquitinated substrate showed that one of the ubiquitin molecules in the chain (the initiator ubiquitin) is unfolded by binding to a groove in Npl4 and insertion of an N-terminal segment into the central pore of the ATPase^4^. Biochemical experiments indicated that no ATP-binding or -hydrolysis is required for the unfolding of the initiator ubiquitin^5^. In the next step, the D2 ring uses the energy of ATP hydrolysis to sequentially move the initiator ubiquitin, all ubiquitin molecules positioned between the initiator and the substrate, and finally the substrate itself through the pore, causing their unfolding. The ubiquitin molecules refold on the trans side, allowing a substrate to rebind to the Cdc48-UN complex, so that the ATPase can act again on a substrate that has already been translocated. How this futile cycle would be disrupted is poorly understood. Furthermore, additional structures are required to confirm the proposed mechanism of substrate processing, particularly because no unfolded ubiquitin was seen in recent structures of the substrate-engaged human p97-UN complex^6,7^.

We have recently shown that a polyubiquitinated substrate can shuttle between the Cdc48/p97 complex and the 26S proteasome before being degraded^8^. Shuttling is facilitated by the proteasome cofactor Rad23 and the Cdc48 cofactor Ubx5, which both reversibly bind folded and unfolded substrates through the substrate-attached ubiquitin chain. Whether other proteins also participate in substrate shuttling between the two molecular machines is unknown, but a possible factor is the deubiquitinating enzyme (DUB) Otu1 (Yod1 in mammals), which binds through its UBX-like (UBXL) domain to Cdc48/p97. The function of Otu1/Yod1 has remained unclear. It was initially proposed that the deubiquitinating activity of Otu1 is required for polypeptide translocation through the central pore^9^. However, subsequent work demonstrated that the removal of the ubiquitin chain is not required^10^. An alternative model was therefore proposed, according to which Otu1 promotes substrate release from the Cdc48 ATPase, thereby breaking the futile cycle of repeated substrate unfolding and facilitating the transfer to the proteasome^10^.

Here, we provide evidence for this model. Using a reconstituted degradation system with purified proteins, we show that Otu1 releases polyubiquitinated substrates from the Cdc48 ATPase by trimming the ubiquitin chain, allowing transfer to the 26S proteasome and subsequent degradation. A cryo-EM structure of human homologs of the Cdc48 complex reveals that Yod1 stabilizes the p97-UN-substrate complex. As in the yeast system, one of the ubiquitin molecules in the chain is unfolded. This initiator ubiquitin molecule is bound to a groove in Npl4 and projects its N-terminus across both ATPase rings, engaging the pore residues of the D2 ring. Thus, the initiation of substrate translocation through the ATPase pore is evolutionarily conserved. Given that p97 is a promising cancer drug target, the structure may also allow the development of specific compounds that interfere with the formation of the ATPase complex.

## Results

### Otu1 promotes substrate transfer from the Cdc48 complex to the 26S proteasome

We first tested the effect of Otu1 on protein degradation using our previously developed *in vitro* degradation assay that utilizes purified components^8^. We initially used as a substrate Ub(n)-TAIL, a purified, polyubiquitinated protein that is labeled with a fluorescent dye. It contains flexible N- and C-terminal segments that enable its direct degradation by the 26S proteasome. When Ub(n)-TAIL was incubated with purified 26S proteasomes, the substrate was degraded to low-molecular weight peptides (**Figure 1A**, lane 2 versus 1). The addition of increasing concentrations of purified Otu1 led to deubiquitination of Ub(n)-TAIL, as seen from the appearance of labeled bands of increased mobility in SDS-PAGE (lanes 3-6). Interestingly, the efficiency of substrate degradation was not affected (**Figure 1A**, lanes 3-6 versus 2), suggesting that Otu1 does not remove the entire ubiquitin chain. This conclusion is in agreement with previous experiments showing that Otu1 releases substrate molecules with chains containing up to ten ubiquitin molecules^9^. Our results show that the remaining ubiquitin chain is sufficient for recognition by the proteasome.

**Fig. 1.**
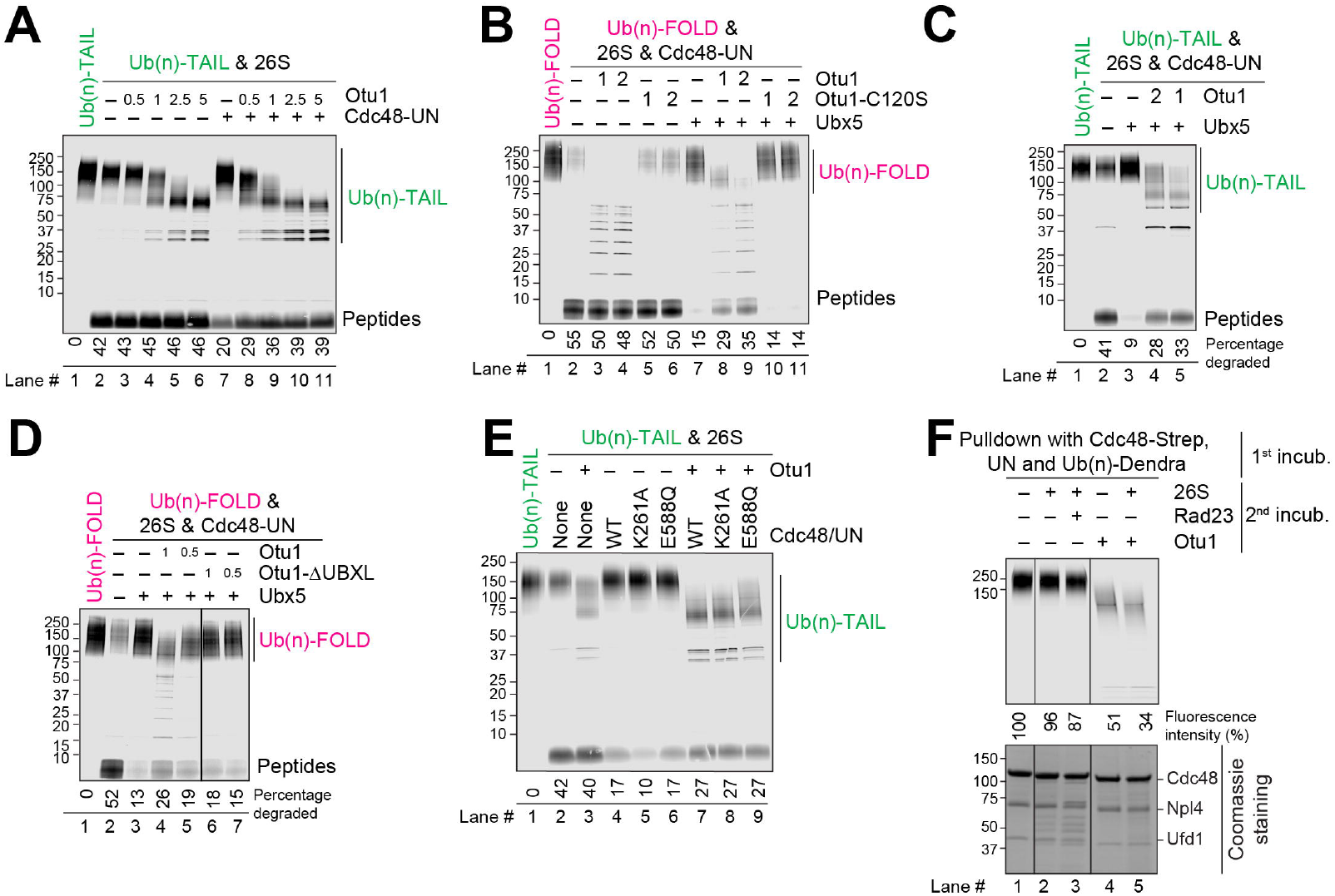
Otu1 stimulates substrate transfer from the Cdc48 ATPase to the 26S proteasome. (**A**) Fluorescently labeled Ub(n)-TAIL was incubated for 60 min with 26S proteasomes. Where indicated, Cdc48-UN and different concentrations of Otu1 (given relative to that of proteasomes) were added. The samples were analyzed by SDS-PAGE, followed by fluorescence scanning. The percentage of substrate degraded was quantified by determining the fluorescence in peptides (after background subtraction) and comparing it to the total fluorescence in each lane of the SDS gel. (**B**) As in (A), but with Ub(n)-FOLD and incubation with both proteasomes and Cdc48-UN. Where indicated, Otu1, an enzymatically inactive Otu1 mutant (Otu1-C120S), or Ubx5 were added. (**C**) As in (B), but with Ub(n)-TAIL. (**D**) As in (B), but with the addition of either full-length Otu1 or Otu1-ΔUBXL. (**E**) As in (A), but with Ub(n)-TAIL and either wild-type (WT) Cdc48 or the indicated mutants. Otu1 was added at the same concentration as proteasomes. (**F**) Fluorescently labeled Ub(n)-TAIL was incubated in the presence of ATP with beads containing streptavidin (Strep)-tagged Cdc48-UN complex (1st incub.). The beads were washed and were then incubated with 26S proteasomes and Rad23 or Otu1, as indicated (2nd incub.). The bound material remaining on the beads was analyzed by SDS-PAGE, followed by fluorescence scanning (upper panel) and Coomassie-blue staining (lower panel). Bound substrate was quantified relative to the material bound in the presence of buffer (lane 1 set to 100%).

The addition of the Cdc48-UN ATPase complex inhibited the degradation of Ub(n)-TAIL (**Figure 1A**, lane 7 versus 2), consistent with Cdc48-UN competing with the 26S proteasome for binding to polyubiquitinated proteins^8^. Importantly, Otu1 was able to relieve this inhibition (**Figure 1A**, lanes 8-11 versus 7), suggesting that Otu1 can release substrates that are trapped on the Cdc48-UN ATPase, thereby allowing their degradation by the proteasome. Otu1 activity is stimulated by the Cdc48 complex, consistent with previous results^9,11^.

We next used as a substrate Ub(n)-FOLD, a polyubiquitinated protein lacking flexible segments. This substrate requires Cdc48-dependent unfolding for proteasomal degradation. When added together with the 26S proteasome and Cdc48-UN, Otu1 caused deubiquitination, but had only a minor effect on the degradation of Ub(n)-FOLD (**Figure 1B**, lanes 3-4 versus 2). These results again suggest that Otu1 does not remove the entire ubiquitin chain, allowing Cdc48-UN and the 26S proteasome to act on the substrate. As expected, the addition of an enzymatically inactive Otu1 mutant (Otu1-C120S) did not cause deubiquitination, but it also had no effect on substrate degradation (lanes 5 and 6). Thus, regardless of whether Otu1 trims the ubiquitin chain, the rebinding of substrate to Cdc48-UN does not effectively compete with substrate transfer to the proteasome.

Next we added Ubx5, a Cdc48 cofactor that helps to recruit polyubiquitinated proteins to Cdc48-UN and therefore counteracts the transfer of substrate to the proteasome^8^. Ubx5 inhibited the degradation of Ub(n)-FOLD (**Figure 1B**, lane 7 versus 2), consistent with the sequestration of the substrate at the ATPase complex. The addition of Otu1 reversed this inhibition in a dose-dependent manner (**Figure 1B**, lane 8 and 9 versus 7), whereas the enzymatically inactive Otu1-C120S mutant was without effect (lanes 10 and 11). Ubx5 also inhibited the degradation of Ub(n)-TAIL when Cdc48-UN was present (**Figure 1C**, lane 3 versus 2), and again this inhibition was alleviated by Otu1 (lanes 4 and 5). These results suggest that trimming of the ubiquitin chain by Otu1 disfavors the rebinding of substrate to the Cdc48-UN-Ubx5 complex and therefore stimulates substrate transfer to the proteasome.

Otu1 contains an UBXL domain that binds to the N domain of Cdc48. Deletion of this domain (Otu1-ΔUBXL) reduced ubiquitin chain cleavage and rendered the protein inactive in stimulating substrate degradation (**Figure 1D**, lane 6 versus 4). Thus, substrate release from Cdc48-UN depends on both Otu1 binding to Cdc48 (**Figure 1D**) and the deubiquitination activity of Otu1 (**Figure 1B**). These results are consistent with previous ones showing that Otu1 has a significantly higher deubiquitination activity when bound to Cdc48 (ref. ^9^).

Next we tested whether Otu1 trims the polyubiquitin chain before substrate translocation through the central pore of Cdc48. To this end, we used Cdc48 mutants defective in ATP hydrolysis and translocation activity, a Walker A mutant in the D1 ring (K261A) and a Walker B mutant in the D2 ring (E588Q)^9^. Wild-type (WT), K261A, and E588Q Cdc48 each inhibited the degradation of Ub(n)-TAIL, with K261A showing the strongest inhibition (**Figure 1E**, lanes 4-6 versus 2). In all cases, Otu1 relieved the inhibition caused by the Cdc48 variants (**Figure 1E**, lanes 7-9 versus 4-6). These results suggest that ubiquitin chain trimming by Otu1 and substrate transfer from Cdc48 to the proteasome can occur prior to initiation of substrate translocation through the ATPase rings. Otu1 stimulates substrate dissociation from Cdc48, which otherwise is slow^8^.

Pull-down experiments confirmed this conclusion. A complex of Cdc48-UN and polyubiquitinated substrates was assembled on beads. After a washing step, the beads were incubated with different combinations of 26S proteasomes, Rad23, and Otu1. 26S proteasomes with or without Rad23 did not efficiently release ubiquitinated substrates from Cdc48-UN, as shown by the large amount of substrate remaining on the beads after the second incubation (**Figure 1G**, lanes 2-3 versus 1). However, the addition of Otu1 caused much of the substrate to be released (**Figure 1G**, lanes 4-5 versus 1). The additional presence of proteasomes had only a small effect (lane 5 versus 4). These results indicate that Otu1-mediated deubiquitination directly leads to substrate release from the Cdc48 ATPase.

### Cdc48 can simultaneously bind several cofactors

Substrate release by Otu1 implies that it can bind to a Cdc48 complex that also contains Ufd1, Npl4, and Ubx5. To investigate whether all four proteins can bind simultaneously, we performed pull-down experiments with a FLAG-tagged version of a catalytically inactive Otu1 mutant (Otu1-C120S-FLAG), which prevents cleavage of the polyubiquitinated substrate. The experiments were performed in the presence of the ATP-analog ATPγS to block Cdc48 function. The samples were subjected to immunoprecipitation with anti-FLAG antibodies and analyzed by SDS-PAGE and fluorescence scanning to detect substrate, and by Coomassie blue staining to identify Cdc48 and the cofactors (**Figure 2A**, upper and lower panels, respectively). The results show that Otu1 alone interacts only weakly with polyubiquitinated substrate (**Figure 2A**, lane 1; upper panel). Substrate binding is drastically enhanced when Cdc48-UN is added (lane 3 versus 1, upper panel). Coomassie-blue staining (lower panel) showed that all components were bound. Importantly, Ubx5 did not bind to the Cdc48-UN-Otu1 complex unless polyubiquitinated substrate was added (lane 5 versus 4). These results demonstrate that Cdc48, Ufd1, Npl4, Otu1, Ubx5, and a polyubiquitinated substrate can assemble into a complex. The cofactors Npl4, Otu1, and Ubx5 all contain a UBX or UBX-like domain that is essential for their interaction with Cdc48 (^8^,^9^,^12^; shown for Otu1 in **Figure 2B**). Each of these domains can interact with one of the six N domains of the Cdc48 hexamer, explaining the simultaneous binding of these cofactors.

**Fig. 2.**
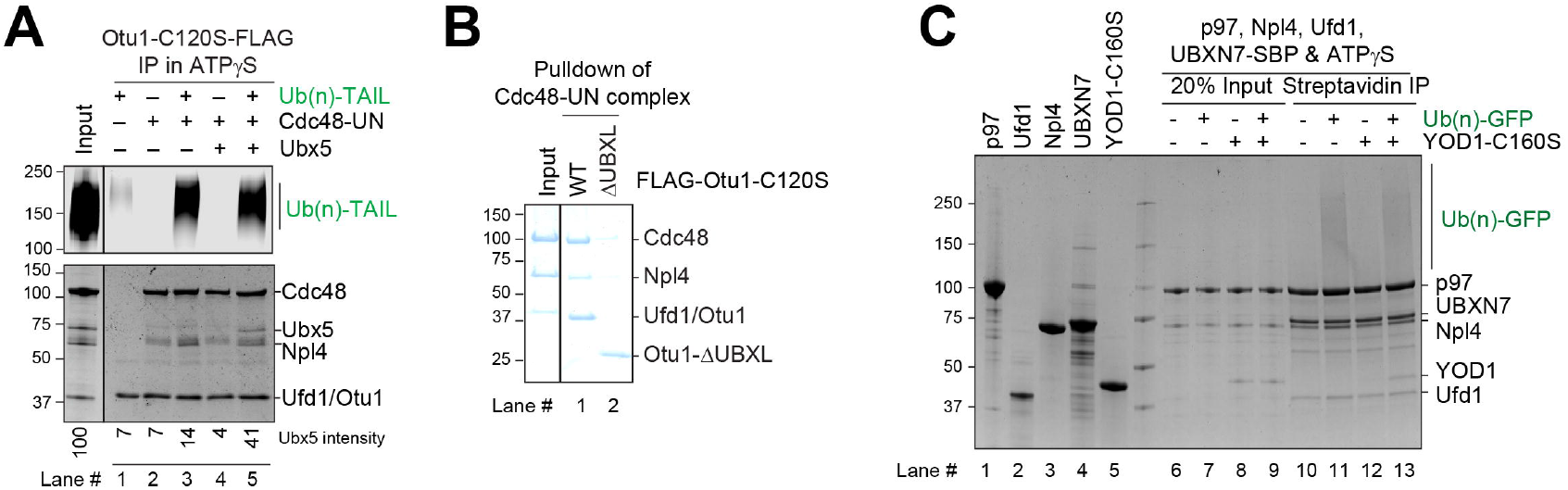
Otu1 binds to the Cdc48 ATPase simultaneously with other cofactors. (**A**) Cdc48-UN was incubated in the presence of ATP*γ*S with different combinations of Ub(n)-TAIL, Otu1-C120S-FLAG, and Ubx5, as indicated. The samples were incubated with beads containing anti-FLAG antibodies and the bound material analyzed by SDS-PAGE, followed by fluorescence scanning (upper panel) and Coomassie-blue staining (lower panel). (**B**) Cdc48-UN was incubated with wild-type Otu1-C120S-FLAG or a mutant lacking the UBXL domain. The samples were incubated with beads containing anti-FLAG antibodies and the bound material analyzed by SDS-PAGE, followed by Coomassie-blue staining. (**C**) Human p97 ATPase, human UN complex, and UBXN7 (Ubx5 homolog) were mixed and incubated with Yod1-C160S (Otu1 homolog) and Ub(n)-TAIL substrate, as indicated. The samples were incubated with streptavidin beads, and the bound material analyzed by SDS-PAGE followed by Coomassie-blue staining. Lanes 1-5 shows the purity of the components.

To test whether a similar supercomplex can form with the human components, we purified p97, the homolog of Cdc48, human Ufd1 (hUfd1), human Npl4 (hNpl4), UBXN7, the homolog of Ubx5, and Yod1, the homolog of Otu1 (**Figure 2C**; lanes 1-5). Pull-down experiments were performed in the presence of ATPγS with an SBP-tagged version of UBXN7, using again an enzymatically inactive mutant of the DUB (Yod1-C160S). After incubation with streptavidin beads, the bound material was eluted with biotin and analyzed by Coomassie blue staining (**Figure 2C**). As with the yeast components, a complex containing all proteins was formed in the presence of polyubiquitinated substrate (**Figure 2C**, lane 13).

### A cryo-EM structure of the substrate-engaged human p97 complex

We used the human p97 complex to determine a cryo-EM structure, hoping that all components would be visible. For structure determination we used a p97 mutant with a Walker B mutation in D2 (E578Q) to slow substrate translocation and the catalytically inactive Yod1-C160S mutant to prevent deubiquitination. Approximately 800,000 particles were subjected to heterogeneous refinement, 3D classification, and non-uniform refinement, resulting in an overall density map of 2.97 Å resolution (**figure S1**). The map was locally refined to 3.05 Å for the D1 ring and the tower formed by hNpl4. Models were built for the ATPase subunits of the D1 and D2 rings and for the hNpl4 tower (**Figure 3A**).

**Fig. 3.**
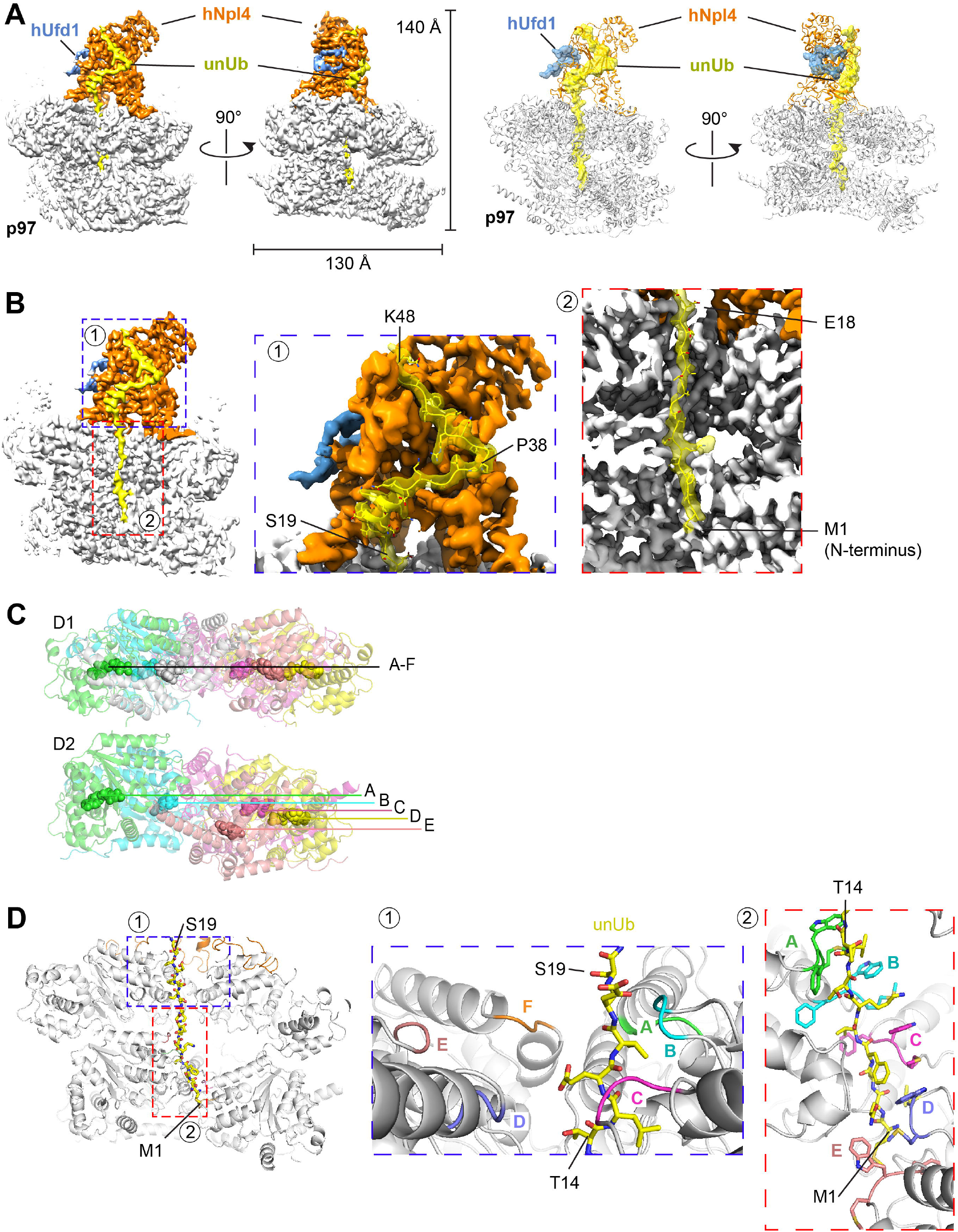
Cryo-EM structure of substrate-engaged p97 ATPase with cofactors. (**A**) A complex containing human p97 ATPase, hNpl4/hUfd1, UBXN7, Yod1, and poly-ubiquitinated substrate (Ub(n)-TAIL) was purified and subjected to cryo-EM single-particle analysis. The left two panels show the density map of the best particle class in two different side views. p97 is shown in grey, hUfd1 in blue, Npl4 in orange, and the unfolded ubiquitin molecule (unUb) in yellow. The right two panels show the corresponding models as cartoons. Ufd1 and unUb are shown as space-filling models. (**B**) The regions indicated as 1 and 2 in the left panel are magnified in the two right panels to show the density of unUb in the Npl4 groove (1) and the central pore of p97 (2). Some residues of unUb are labeled. (**C**) Arrangements of the six protomers (A to F) in the D1 and D2 rings. The positions of the nucleotide in each protomer are labeled. Note that D1 ring is almost planar, while the protomers of the D2 ring are arranged as a staircase. (**D**) Boxed areas 1 and 2 in the left panel are magnified and the corresponding models are shown on the left. ubUb is shown as a stick model. The pore loops of the D1 ring (inset #1) are shown in different colors. The aromatic pore loop residues in the D2 ring (inset #2) are shown as stick models.

hNpl4 is anchored to the D1 ATPase ring by its two Zn^2+^-finger domains (ZF-1 and ZF-2), a β-strand finger, and an N-terminal bundle (NTB) domain (**figure S2A**). These interactions are similar to those of yeast Npl4 (ref.^4^), except that the NTB in mammals is significantly shorter (**figure S2B**) and binds to D1 closer to the central pore (**figures S2A** and **S2C**). In our structure the hNpl4 tower is fixed in a defined position and both Zn^2+^-fingers interact with D1, whereas other authors reported that the hNpl4 tower was tilted to different degrees and ZF-2 was not always engaged^7^. The differences might be due to the presence of an unfolded ubiquitin molecule and/or the additional presence of Yod1 and UBXN7 in our structure.

Density for an unfolded ubiquitin molecule (unUb) was seen from the groove in Npl4 all the way through both ATPase rings (**Figure 3A, 3B**). As in the yeast Cdc48 ATPase complex^4^, the Npl4 binding groove that has a prominent kink, dividing the groove into upper and lower parts (**Figure 3B**). The upper part accommodates residues P38 to K48 of unUB in an unfolded conformation, whereas the lower part binds residues S19 to P38, which includes the single helix of ubiquitin (residues I23 to Q31) that remains folded (**Figure 3B**). Because the resolution of the mammalian complex is significantly higher than that of the yeast complex (∼3Å versus 4Å), many amino acid side chains could be modeled. Most unUb-interacting residues in the hNpl4 groove are conserved, as noticed before (ref). However, two differences to the yeast complex were observed (**figure S2D**). First, F45 of unUb intercalates between F382 and H231, whereas this residue does not contribute to the interaction in yeast Npl4. Second, K33 and R42 of unUb bind to an acidic patch that includes E383 and D451, which is only present in the groove of human Npl4. Binding to unUb also leads to a conformational change of the Npl4 groove. In the isolated human UN complex, the lower groove of hNpl4 is occluded^13^; in our active complex, two helices shift outwards, widening the groove so that it can accommodate unUb (**figure S2E**).

The N-terminus of unUb extends through the central pore of p97 all the way to the D2 ring (**Figure 3B**). As in the yeast complex, the D1 ring pore loops adopt a planar arrangement (**Figure 3C**), with the loops of subunits A, B, and C contacting the unfolded ubiquitin chain (**Figure 3D; inset #1**). The D2 pore loops form a staircase, with subunit A on top and subunit E on the bottom (**Figure 3D; inset #2**). Five of the loops contact the polypeptide. This arrangement supports a model in which sequential ATP hydrolysis by the D2 subunits allows the pore loops to function as a conveyer belt. In this model, subunit A binds to the top of the polypeptide chain and the subunits then move downwards, dragging the polypeptide chain with them. At the lowest position, subunit F disengages from the substrate and becomes flexible (and invisible in our structures); it can then move to the top position and start a new cycle.

Density for the N domains of p97 and for the UBXL domain of Npl4 was weak, but after local classification and refinement, a particle class was obtained in which three N domains were much more visible than the other three. The visible N domains are associated with subunits A, C, and F of the hexameric ring (**Figure 4A**). This class also displayed density for the Npl4 tower and for unUb in the central pore and it showed the D2 ring in the staircase conformation characteristic of substrate engagement (**Figure 4A**). The three visible N domains are in the up-conformation that likely requires the associated D1 subunits to be in the ATP-bound state^9^. Density associated with one of the N domains (of subunit C) was visible, but did not allow an unambiguous assignment to one of the three UBX-like domains (those of Npl4, Yod1, and UBXN7). However, the density most likely belongs to UBXL domain of Npl4, because an AlphaFold^14^-model docked into it places the C-terminus the domain only 20 amino acids from the Npl4 tower (**figure S3A**; **Figure 4A**). No density for the other two UBX or UBX-like domains was visible, perhaps because the N domains are too flexible, but it is tempting to speculate that they are bound to the visible N domains associated with subunits A and F. Although UBX domains can bind to both the up- and down-conformations of N domains^15,16^, they might stabilize the up-conformation.

**Fig. 4.**
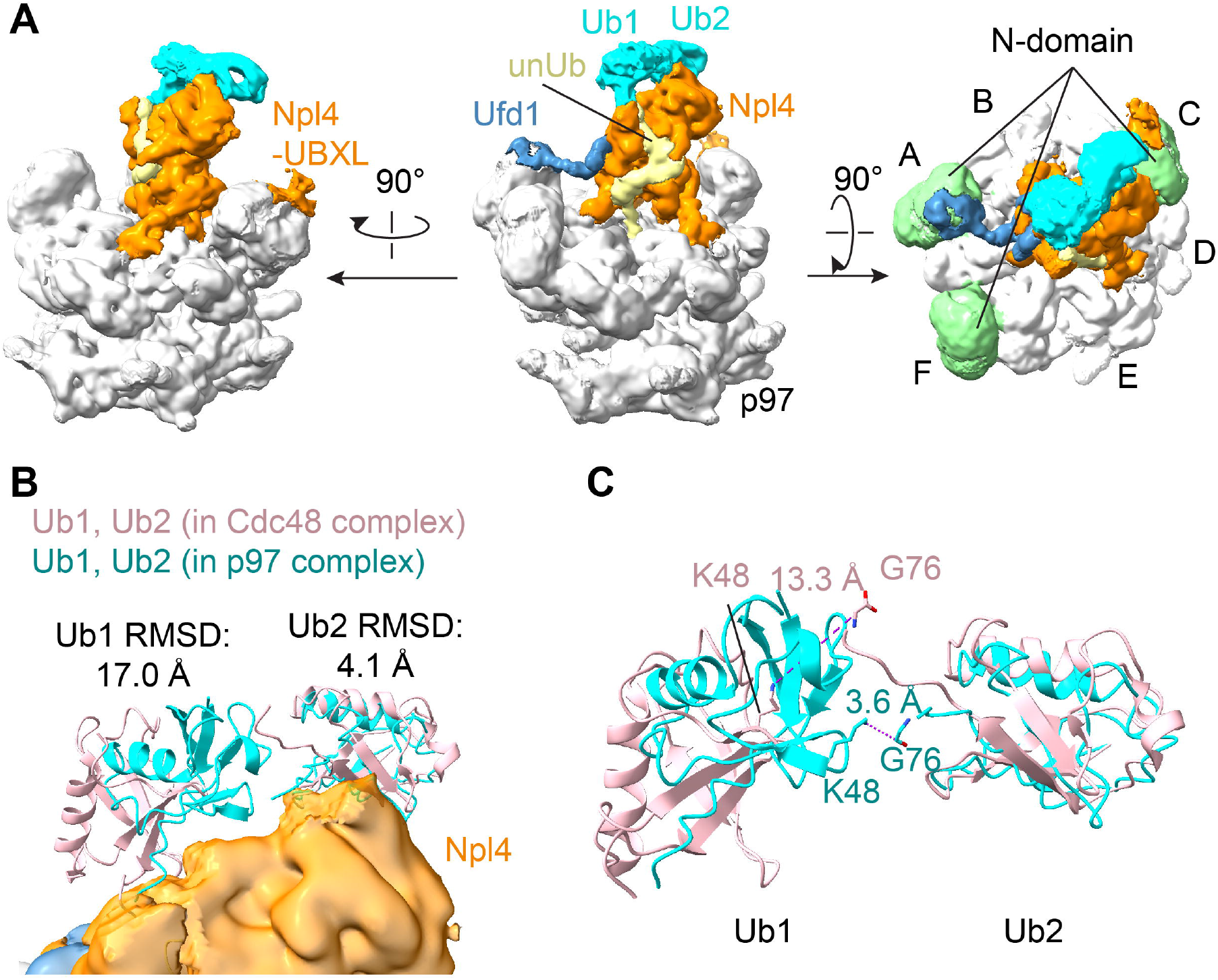
Interactions of the N domain of p97. (**A**) The cryo-EM map was locally refined based on a mask containing the Npl4 tower, the D1 ring, and three N domains. The composite map of the locally refined map and the overall map are shown in three different views. p97 is shown in grey, Ufd1 in blue, Npl4 in orange, the unfolded ubiquitin molecule (unUb) in yellow, and the following two folded ubiquitin molecules (Ub1 and Ub2) in cyan. In the right most panel, the visible three N domains are shown in green. (**B**) Models for Ub1 and Ub2 are shown as cyan ribbons. The corresponding ubiquitin molecules of the yeast complex are shown as pink ribbons, based on an alignment of Cdc48-UN (PDB: 6OA9)^4^ with p97-UN. RMSD values were calculated for each pair of ubiquitin molecules. (**C**) Magnified view of the arrangement of Ub1 and Ub2. Note that in the human complex (cyan) K48 of Ub1 is close to G76 of Ub2, whereas in the yeast complex (pink) the distance is too large for the two ubiquitin molecules to be linked (dashed lines with labeled distance).

Density for two folded ubiquitin molecules (Ub1 and Ub2) was visible on the top of the Npl4 tower (**Figure 4A;B**), as in previous structures obtained with polyubiquitinated substrate^4^. These ubiquitin molecules follow unUb in the chain (distal ubiquitin molecules). Interestingly, Ub2 was approximately at the same position as in the yeast complex, but Ub1 was much closer to Ub2 (**Figure 4B**)^4^. In fact, in the yeast structure, the two ubiquitin molecules are too far apart to be linked, suggesting that only Ub2 was actually bound. In contrast, in the human complex K48 of Ub1 is close to the C-terminus of Ub2 (**Figure 4C**), indicating that both Ub1 and Ub2 are bound to the top of the Npl4 tower.

As in the yeast complex^4,17^, only a portion of Ufd1 is visible (the UT6 domain), whereas the ubiquitin-binding UT3 domain is too flexible to visualize. The central part of the UT6 domain forms a loop that associates with Npl4 (**figure S3B**). The loop is slightly different from that seen in the crystal structure of the UN complex in isolation (**figure S3B**)^13^. The loop is preceeded by a SHP segment (SHP1) that likely binds to the N domain of subunit A, as there is density for the linker region (**figure S3C**). The second SHP segment (SHP2) is invisible, as in the yeast structure, but probably interacts with the N domain of subunit F. The exact function of Ufd1 is still unclear, but it is required for substrate translocation^10^.

## Discussion

Here we show that the deubiquitinating enzyme Otu1 trims the ubiquitin chain bound to the Cdc48 ATPase complex, releasing substrates before they are translocated through the central pore of the double-ring ATPase. The released substrate still carries sufficient ubiquitin molecules for degradation by the 26S proteasome. Substrate dissociation from Cdc48-UN and binding to the 26S proteasome are possible because the ATPase complex requires slightly longer polyubiquitin chains than the proteasome (5 ubiquitin molecules versus 3-4; the human p97 prefers even longer chains)^9,18^. The action of Otu1 breaks the futile cycles of substrate translocation through the Cdc48 pore and therefore facilitates substrate transfer from the ATPase to the proteasome. Otu1 belongs to a small group of DUBs that promotes, rather than counteracts, protein degradation. This group includes Rsp11 that removes the ubiquitin chain from substrates prior to their translocation into the proteolytic chamber of the proteasome^19,20^.

Otu1 likely cooperates with Ubx5 and Rad23, the recruitment factors for Cdc48-UN and the 26S proteasome, respectively. Polyubiquitinated substrate bound to the Cdc48-UN-Ubx5 complex can spontaneously dissociate and then be recruited by the 26S proteasome/Rad23 complex. However, when the ubiquitin chain is long, the rebinding of substrate to the ATPase complex is favored, so that only a fraction of the substrate is transferred to the proteasome for degradation. Otu1 reduces the efficiency of rebinding to the ATPase complex and thus increases the efficiency of transfer and degradation. Otu1 is not essential for the viability of *S. cerevisiae* cells^9,11^, so it is either a facilitating factor or only required under certain conditions. For example, Otu1 might reduce the load of the Cdc48 ATPase complex when misfolded polyubiquitinated proteins accumulate that bind to the ATPase but do not require unfolding.

Our model implies that Otu1 can bind to Cdc48 simultaneously with the cofactors Npl4, Ufd1, and Ubx5, as well as polyubiquitinated substrate. This prediction is indeed confirmed by pull-down experiments. Moreover, our experiments suggest that substrate binding promotes the interaction of Cdc48 with the cofactors. This synergistic interaction is also seen with the human proteins and is consistent with earlier results showing that binding of the UN complex stimulates the binding of the human Ubx5 homolog^21^. However, it is likely that the cofactors are not always bound at the same time and could associate with Cdc48/p97 in different combinations. How exactly substrate and the cofactors cooperate requires a structure in which all components are visualized.

An important result of our study is the demonstration that the initiation of substrate processing by the ATPase complex is evolutionary conserved. Our cryo-EM structure of the human complex is quite similar to the one obtained with *S. cerevisiae* components. As in yeast^4^, one of the ubiquitin molecules in the chain (the initiator) is unfolded. This ubiquitin molecule is bound to a groove in Npl4 and projects its N-terminal segment across both ATPase rings. The interaction with the pore loops of the D2 ring indicates that a conveyor belt mechanism is used to pull on the N-terminus of the initiator ubiquitin. It remains unclear why recent cryo-EM structures of Cdc48-UN with SUMO– polyubiquitin hybrid chains and of human p97-UN with polyubiquitinated substrate did not show a ubiquitin segment in the central pore^22^ or unfolded ubiquitin in Npl4 and the pore^6,7^.

Our structure supports a model in which the ATPase first moves the initiator ubiquitin through the central pore, then all ubiquitin molecules positioned between the initiator and the substrate (proximal ubiquitins), and finally the substrate itself. All polypeptides moving through the central pore are unfolded, but the proximal ubiquitin molecules refold when they emerge on the trans side of the double-ring ATPase. The distal ubiquitin molecules are not translocated through the central pore and are therefore never unfolded^5^. They likely move through lateral openings between two protomers of the two ATPase rings.

p97 has long been considered a promising target of cancer drugs^23^. Inhibitors of p97 have entered clinical trials for solid tumors and myeloid leukemia, but toxicity remains an issue^24^. Recent efforts have therefore focused on developing specific inhibitors that disrupt the p97-UN complex. Thonzonium bromide has been shown to inhibit tumor growth by binding to the interface between p97 and Npl4^25^. and peptide inhibitors derived from Ufd1 compete with Npl4 for binding and inhibit the unfolding activity of the p97-UN ATPase^26^. Our structure provides the basis for yet another approach in which drugs would prevent the binding of unfolded ubiquitin to the Npl4 groove.

## METHODS

## RESOURCE AVAILABILITY

### Lead Contact

Further information and requests for resources and reagents should be directed to and will be fulfilled by the Lead Contact, Tom Rapoport.

### Materials Availability

All unique/stable reagents generated in this study are available from the Lead Contact with a completed Materials Transfer Agreement.

## EXPERIMENTAL MODEL AND STUDY PARTICIPANT DETAILS

### Bacterial cultures

Bacterial cultures were handled as described^8^. Briefly, plasmids for protein expression were transformed into *E. coli* BL21 CodonPlus (DE3) RIPL cells (Agilent). Cultures grown in Terrific Broth were induced with 0.1 mM isopropyl β-D-thiogalactopyranoside (IPTG) at an OD600 of 0.8 and then transferred to 16°C for 16 h.

## METHOD DETAILS

### Plasmids

Plasmids were prepared as described^8^. Briefly, wild-type Cdc48/p97 and their variants were cloned into the pET28 vector with an N-terminal His6-tag and a TEV-protease cleavage site. For Cdc48 pull-down experiments, two copies of strep tag (WSHPQFEK) were added at its C-terminus. Ufd1 was cloned into the pK27 vector followed by an N-terminal His14-SUMO (small ubiquitin-like modifier) tag, as described^8^. Npl4 was cloned into the pET21 vector with a C-terminal His6-tag or FLAG-His6 (DYKDDDDKGLEHHHHHH) tag, as described^8^.

Wild-type Otu1/Yod1 and their variants were cloned into the pET28 vector with an N-terminal His6-tag and a TEV-protease cleavage site followed by a FLAG tag. Wild-type Ubx5 or UBXN7 was cloned into the pET28 vector with an N-terminal His6-tag and a C-terminal SBP tag.

A bacterial expression plasmid coding for an N-end degron (NeD) sequence with lysine-less super-folder GFP (NeD-sfGFP) and Ub(n)-FOLD has been described^5,8^.

### Protein purifications

Protein purification was performed as described^8^. Briefly, bacterial cells were harvested by centrifugation at 5,000 x g for 10 min at 4°C and resuspended in cold wash buffer (50 mM Tris-HCl, pH 8, 320 mM NaCl, 5 mM MgCl_2_, 10 mM imidazole, 0.5 mM ATP) supplemented with phenylmethylsulfonyl fluoride (PMSF; 1 mM) and a protease inhibitor cocktail, and lysed by sonication. After ultracentrifugation, the supernatant was incubated with Ni-NTA resin that was pre-equilibrated with wash buffer. The beads were washed three times with wash buffer and protein eluted with elution buffer (50 mM Tris-HCl, pH 8, 150 mM NaCl, 5 mM MgCl_2_, 400 mM imidazole). The eluted protein was concentrated and subjected to size-exclusion chromatography (SEC).

Untagged Ufd1/Npl4 (UN) complex, Ub-G76V-Dendra, and NeD-sfGFP were purified as described^8^. FLAG-26S proteasomes were expressed in yeast cells and purified as described^8,27^. *S. cerevisiae* ubiquitin was purchased from Boston Biochem. Photoconversion, dye labeling, and *in vitro* ubiquitination of Ub-G76V-Dendra (for Ub(n)-FOLD), and dye labeling and *in vitro* ubiquitination of NeD-sfGFP (for Ub(n)-TAIL) have been described^8,10^.

### Substrate degradation assays

Substrate degradation assays were carried out as described^8^. Briefly, 100 nM of fluorescently labeled substrate was mixed with 200 nM 26S proteasomes, Cdc48, UN, Otu1, Rad23, or Ubx5 in 26S reaction buffer (60 mM HEPES, pH 7.4, 20 mM NaCl, 20 mM KCl, 10 mM MgCl_2_, 2.5% glycerol, and 1 mM TCEP) and 0.5 mg/ml protease-free bovine serum albumin (BSA). The reaction was incubated with an ATP regeneration mixture (5 mM ATP, 16 mM creatine phosphate, 0.03 mg/mL creatine kinase) for 1 h. The samples were subjected to SDS-PAGE, followed by fluorescence scanning on an Odyssey imager (LI-COR).

### Pull-down experiments

The two-step pull-down experiments in Figure 1F were performed, as described^8^, with some modifications. Briefly, 5 µl pre-equilibrated FLAG antibody M2 agarose beads (Sigma) were first incubated with 0.2 µM Cdc48, 0.2 µM Ufd1, 0.2 µM Npl4-FLAG, 0.2 µM polyubiquitinated substrate in 26S reaction buffer supplemented with 5 mM ADP for 1 h at 4°C. The beads were washed three times with 26S reaction buffer. The beads were then incubated with 0.2 µM Otu1, 26S proteasomes, or Rad23 in 26S reaction buffer supplemented with ATP for 20 min at 30°C. The beads were washed again and proteins were eluted with 20 µl of 26S reaction buffer supplemented with 0.2 mg/ml 3xFLAG peptide (Bimake). The eluted samples were subjected to SDS-PAGE, followed by fluorescence scanning and Coomassie-blue staining.

The pull-down experiments in Figure 2A were performed as described^8^, with the following modifications. 10 µl pre-equilibrated FLAG antibody M2 agarose beads (Sigma) were incubated with 1 µM Otu1-C120S-FLAG, 1 µM Cdc48, 1 µM UN, 1 µM Ubx5, and 1 µM Ub(n)-TAIL in SEC buffer supplemented with 5 mM ATPγS for 1 h at 4°C. The beads were washed three times with SEC buffer and bead-bound proteins were eluted with 20 µl of SEC buffer supplemented with 0.2 mg/ml 3xFLAG peptide (Bimake).

For the pull-down experiments in Figure 2B, a mixture of 1 uM Cdc48 and 1 uM UN in 5 mM ADP was mixed and incubated for 0.5 h at 4°C. 10 µl pre-equilibrated FLAG antibody M2 agarose beads (Sigma) were incubated with 1 µM Otu1-C120S or Otu1-ΔUBXL-C120S in SEC buffer for 1 h at 4°C. The beads were washed three times with SEC buffer followed by incubation with the pre-assembled Cdc48-UN complex for 1 h at 4°C. The beads were washed three times with SEC buffer and bead-bound proteins were eluted with 20 µl of SEC buffer supplemented with 0.2 mg/ml 3xFLAG peptide (Bimake). The eluted samples were subjected to SDS-PAGE, followed by Coomassie-blue staining.

For the pull-down experiments in Figure 2C, 10 µl pre-equilibrated streptavidin agarose beads (Thermo Fisher) were incubated with 1 µM UBXN7-SBP, 1 µM p97, 1 µM human UN, 1 µM Yod1-C160S, and 1 µM Ub(n)-TAIL in SEC buffer supplemented with 5 mM ATPγS for 1 h at 4°C. The beads were washed three times with SEC buffer and bead-bound proteins were eluted with 20 µl of SEC buffer supplemented with 2 mM biotin.

### Cryo-EM sample preparation and data collection

100 µl pre-equilibrated streptavidin agarose beads (Thermo Fisher) were incubated with 1 mL mixture of 0.2 µM UBXN7-SBP, 0.2 µM p97-E578Q, 0.2 µM human UN, 0.2 µM Yod1-C160S, and 0.2 µM Ub(n)-TAIL in SEC buffer supplemented with 5 mM ATP for 1.5 h at 4°C. The beads were washed three times with SEC buffer supplemented with ATP, and bead-bound proteins were eluted with 300 µl of SEC buffer supplemented with 2 mM biotin and 5 mM ATP. The eluted proteins were concentrated to 30 µl and incubated with 0.05% glutaraldehyde for 30 min at room temperature. The reaction was quenched with 40 mM Tris pH 7.5.

All samples for cryo-EM were prepared with a Thermo Fisher Vitrobot Mark IV system, maintained at 100% humidity at 4°C. Blot times of 5.0 s were used 30s after 3 μl sample application to Quantifoil Au 1.2/1.3 200 mesh grids that had previously been plasma-treated in a Pelco easiGlow at 0.39 mBar, 15 mA for 60 s.

6,426 micrographs were collected on a Thermo Fisher Titan Krios operating at 300 kV, with GIF BioQuantum energy filter and a Gatan K3 camera in super resolution mode (super resolution pixel size of 0.4135 Å/pixel). The total exposure time of 10 s was divided into 50 frames (total dose of approximately 50 (e−/Å^2^), with a defocus range of −0.8 μm to −2.0 μm. The objective and C2 apertures were set to 70 μm.

### Image processing

All subsequent image processing steps were carried out using Cryosparc^28^. Frame alignment and dose-weighting was carried out using Patch Motion Correction. CTF correction was performed with Patch CTF. Density visualization was performed using UCSF ChimeraX^29^.

After picking a total of 2,044,375 particles, an initial round of 2D classification resulted in 801,542 particles, which were subjected to heterogeneous refinement. One class of 361,890 particles was selected for further analysis. The particles were sub-classified, resulting in two classes totaling 113,094 particles. This class was then sub-classified into two classes, one of which was chosen for further processing based on the density for Npl4. The resulting 65,959 particles were subjected to refinement and gave rise to a map of 2.97 Å resolution. To better visualize the Npl4 tower, the initial model was locally refined with a mask containing the p97 D1 ring and the Npl4 tower, resulting in a 3.05 Å map, where the D2 ring density was significantly weaker in local resolution compared to the rest of the density.

### Model Building

Initial models were built into the map using existing structures of p97 (PDB: 7LMZ)^6^, human Ufd1 (PDB: 7WWQ)^13^, human Npl4 (AlphaFold Database: AF-Q8TAT6-F1)^30^, and unfolded ubiquitin (PDB: 6OA9)^4^ with rigid body fitting. Model building was then carried out in Coot. Amino acid side chain positions were modified based on the density.

For the model shown in Figure 4 and figure S3, the structure of ubiquitin molecules (PDB: 6OA9)^4^ was fit as a rigid-body into the density. The structures of p97 N domains (PDB: 7R7S)^16^ were fit as rigid-bodies into the density of subunits A, C, and F. The structure of human Ufd1 SHP1 motif was modeled after a SHP1-N domain structure (PDB: 5B6C)^31^.

## QUANTIFICATION AND STATISTICAL ANALYSIS

Quantification of bands after fluorescence scanning of gels was carried out using the ImageStudio software (LI-COR) as described^8^.

## ACKNOWLEDGEMENTS

We thank R. Yan at the HHMI Janelia Research Campus cryo-EM facility for assistance with microscope operation and data collection, Z. Li, R. Walsh, C. Leistner, M. Mayer, R. Nair, and S. Rawson at the Harvard cryo-EM Center for Structural Biology for assistance with sample preparation, microscope operation, data collection, and image processing, the SBGrid team for software and workstation support. This work was supported by a NIGMS grant (R01 GM052586) to T.A.R. T.A.R. is a Howard Hughes Medical Institute Investigator.

## AUTHOR CONTRIBUTIONS

H.L. performed all experiments, H.G. helped with cryo-EM sample preparation and data processing, T.A.R. supervised the project. H.L. and T.A.R. wrote the manuscript.

## DATA AVAILABILITY STATEMENT

Cryo-EM maps have been deposited in the Electron Microscopy Data Bank (EMDB, www.ebi.ac.uk/pdbe/emdb/) under the accession codes EMD-73365 (composite map), EMD-73375 (globally refined map), and EMD-73376 (map locally refined on p97 D1 and Npl4). The atomic model has been deposited in the Protein Data Bank under the accession codes PDB 9YRC.

