## Supplemental material for "The deubiquitinating enzyme Otu1 releases substrates from the conserved initiation complex of the Cdc48/p97 ATPase for proteasomal degradation"

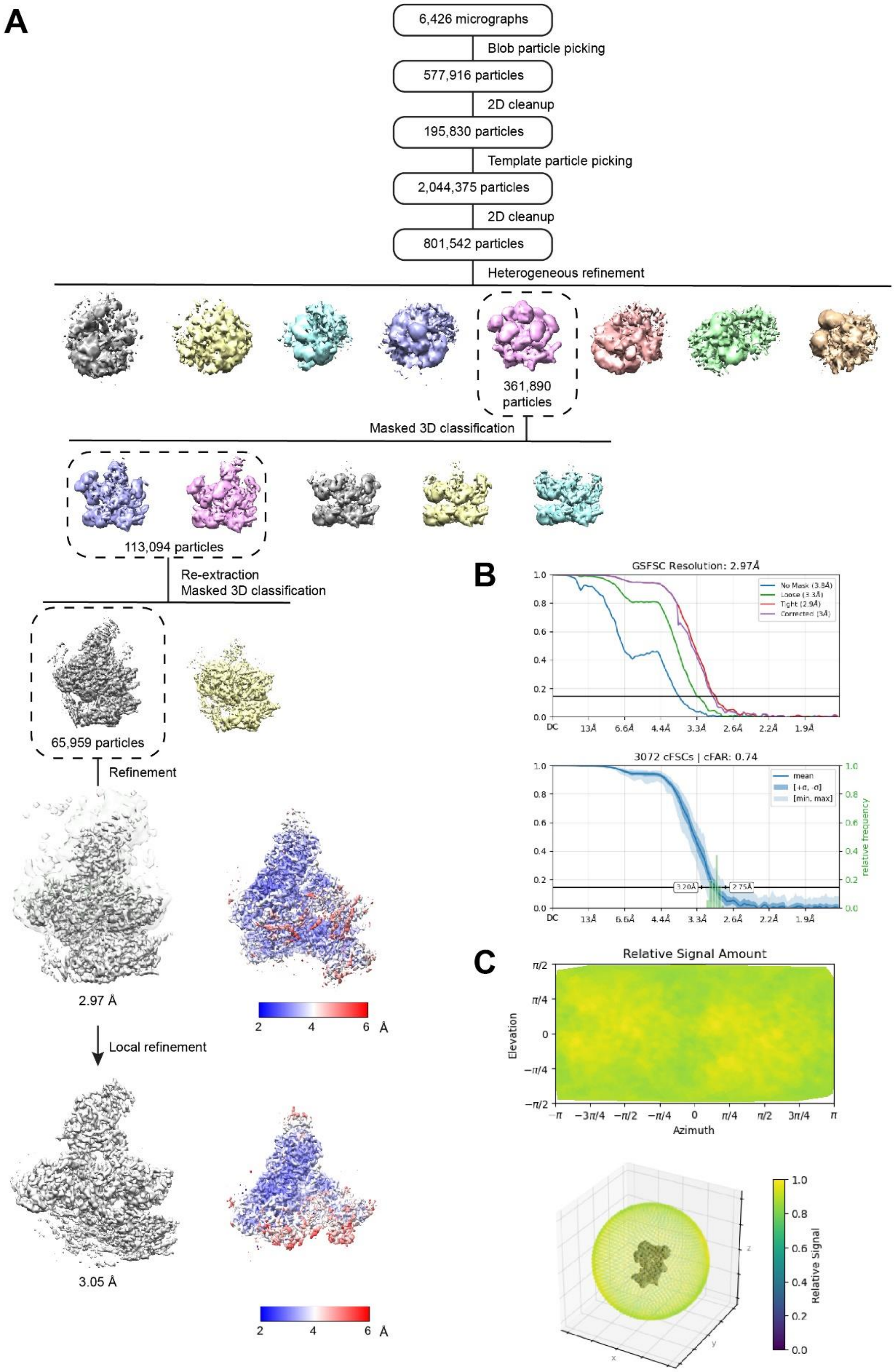

**Fig. S1. Cryo-EM analysis of the substrate-engaged p97 complex.**

(**A**) Image processing workflow for 3D classification and refinement. Particle classes after heterogeneous refinement are shown as side views in different colors. The indicated class was used for masked 3D classification. Two classes were combined and subjected to further masked classification. The indicated class was then subjected to refinement, resulting in a reconstruction that is shown in two views, with the local resolution shown on the right (scale below). Local refinement resulted in the reconstruction shown on the bottom.

(**B**) Up – Gold Standard Fourier Shell Correlation (GSFSC) curves with indicated resolution at FSC = 0.143. Down – conical FSC (cFSC) summary plot, including the cFAR (conical FSC Area Ratio) score[20].

(**C**) Relative signal visualized in a 2D azimuth-elevation chart (up), and in a 3D-colored scatter plot (down) with a low-pass-filtered volume embedded within. Low relative signal indicates regions with under-represented views (visualized by dark blue color). High relative signal is indicated by green and yellow colors.

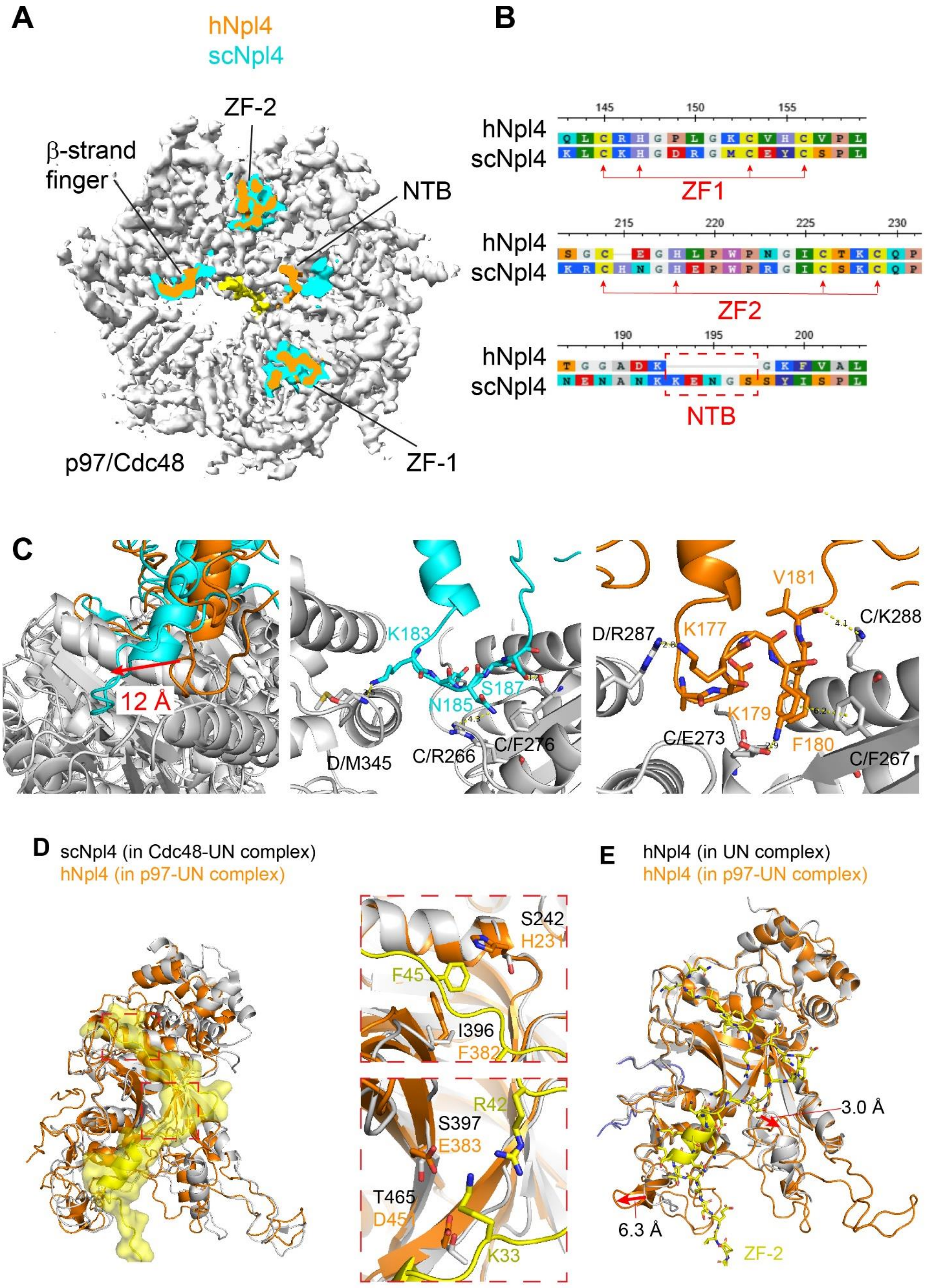
A
hNpl4
scNpl4
ZF-2
β-strand finger
NTB
p97/Cdc48
ZF-1
B
hNpl4
scNpl4
ZF1
ZF2
NTB
C
12 Å
K183
N185
S187
D/M345
C/R266
C/F276
V181
K177
D/R287
C/K288
K179
C/E273
F180
C/F267
D
scNpl4 (in Cdc48-UN complex)
hNpl4 (in p97-UN complex)
S242
H231
F45
I396
F382
R42
S397
E383
T465
K33
E
hNpl4 (in UN complex)
hNpl4 (in p97-UN complex)
3.0 Å
6.3 Å
ZF-2

**Fig. S2. Comparison of the human and yeast Npl4 towers.**

(**A**) Top view of the density map of the substrate-engaged human p97-UN complex, cut close to the surface of the D1 ring. Density for Npl4 is shown in orange. For comparison, the density of yeast Npl4 in the Cdc48-UN complex is shown in cyan. Note that the positions of ZF-1, ZF-2, and the β-strand finger positions are very similar, while there are differences in the binding of the NTB.

(**B**) Sequence alignment of human and yeast Npl4 regions. Note that the NTB is shorter in human Npl4.

(**C**) Magnified views of the interaction between the NTB and Cdc48/p97. The left panel shows an overlay, and the right panels show the interactions separately, highlighting the distinct modes of binding.

(**D**) The left panel shows an overlay of the human (orange) and yeast (gray) Npl4 towers as cartoons with the unfolded ubiquitin molecule (unUb) as a space-filling model in yellow. The right panels show interactions between human Npl4 and unUb not seen in the yeast complex.

(**E**) Comparison of the Npl4 tower in the substrate-engaged p97-UN complex (in orange) with the tower in the isolated Ufd1/Npl4 complex (PDB: 7WWQ)[10] (in yellow). Note that the Npl4 groove widens to accommodate unUb.

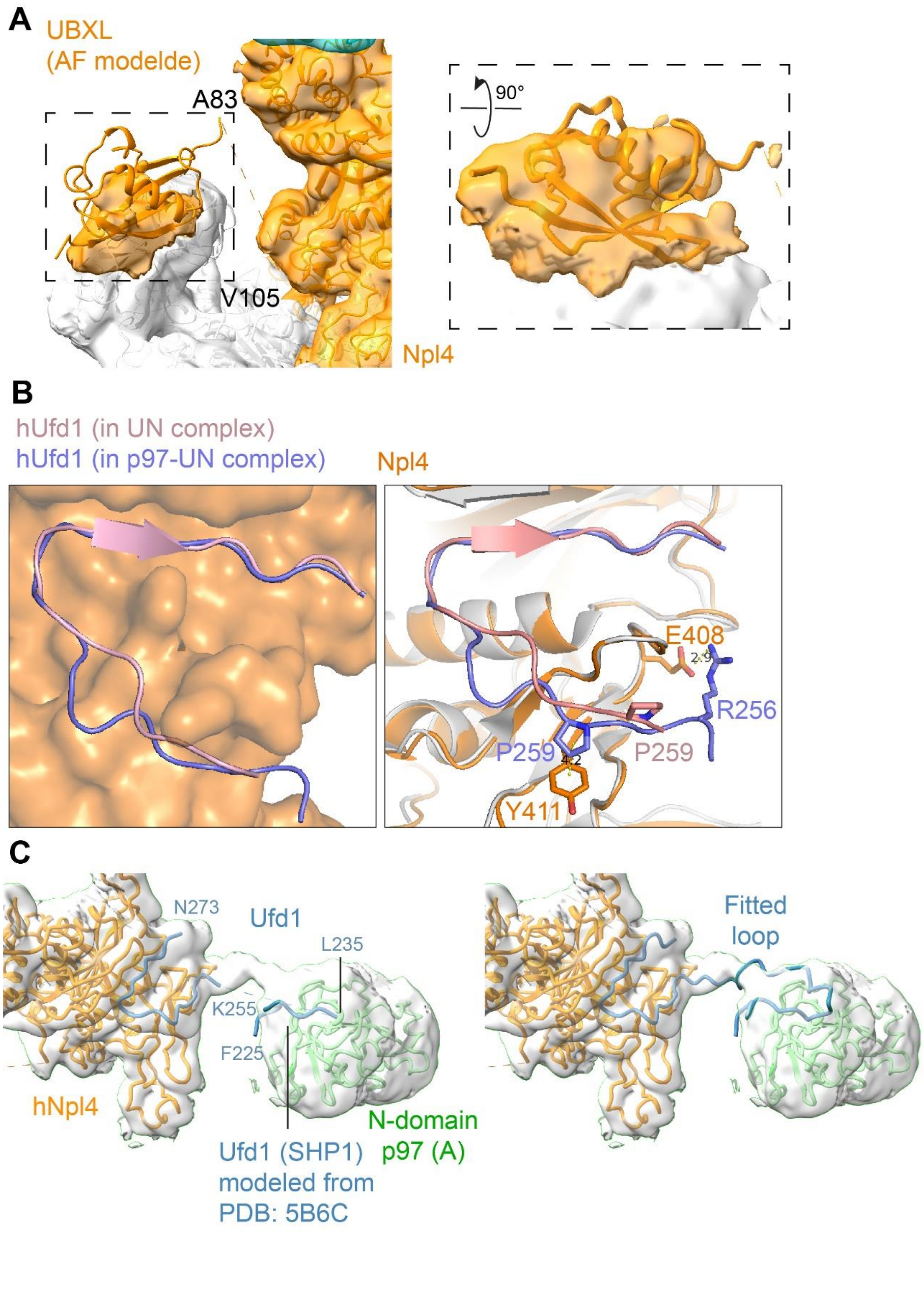

A
UBXL
(AF modelde)
A83
V105
Npl4
90°
B
hUfd1 (in UN complex)
hUfd1 (in p97-UN complex)
Npl4
E408
2.9
R256
P259
P259
4.2
Y411
C
N273
Ufd1
L235
K255
F225
hNpl4
Ufd1 (SHP1)
modeled from
PDB: 5B6C
N-domain
p97 (A)
Fitted
loop

**Fig. S3. Interactions of Ufd1 and Npl4 with p97.**

(**A**) The left panel shows the magnified view of the UBXL domain of Npl4. The structure of Npl4 UBXL bound to the N domain of p97 was predicted by AlphaFold3 and docked into the composite density map. The invisible 23 residues between the Npl4 tower and the UBXL domain are indicated as a dashed line. The right panel shows the further magnified view of UBXL domain fitted into density contoured at 0.02.

(**B**) The left panel shows the human Ufd1 loop (blue ribbon) bound to Npl4 in the substrate-engaged p97-UN complex and in the isolated Ufd1/Npl4 complex (PDB: 7WWQ)[10] (pink ribbon). Npl4 is shown as a space-filling model. The right panel shows differences between the two structures.

(**C**) The left panel shows the semi-transparent density map in grey and models for Npl4 (in orange) and two Ufd1 segments (in blue). The interaction between one of the segments (Shp1 segment) with the N domain is based on a crystal structure (PDB: 5B6C)[21]. The right panel shows a model in which the two Ufd1 fragments are connected with a segment that fits into the density map.

**Raw images of all gels**

**Figure 1A**

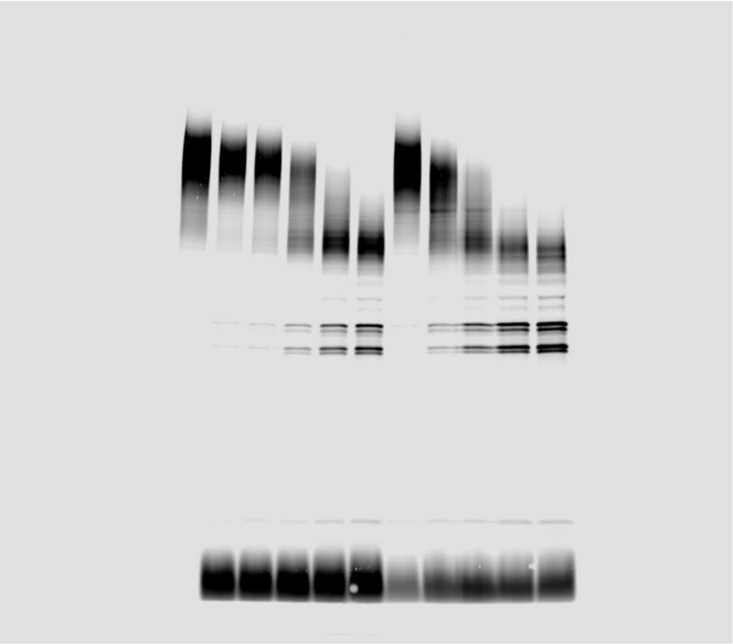

Figure 1B

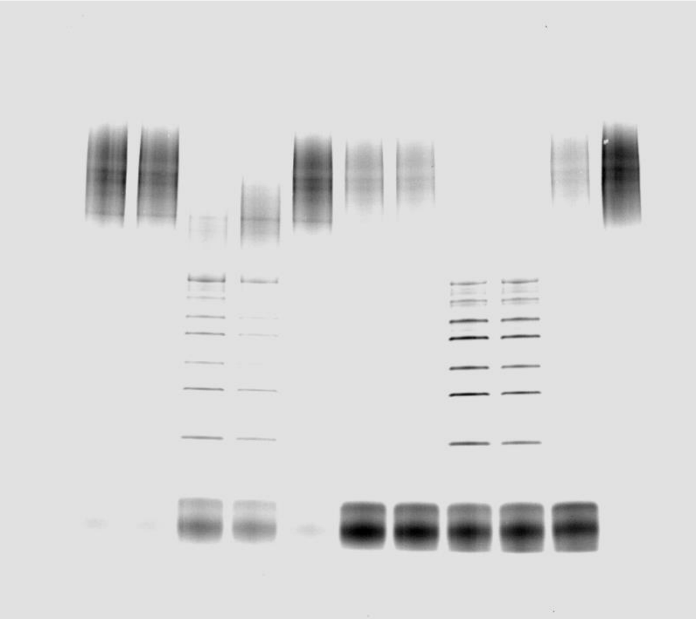

Figure 1C

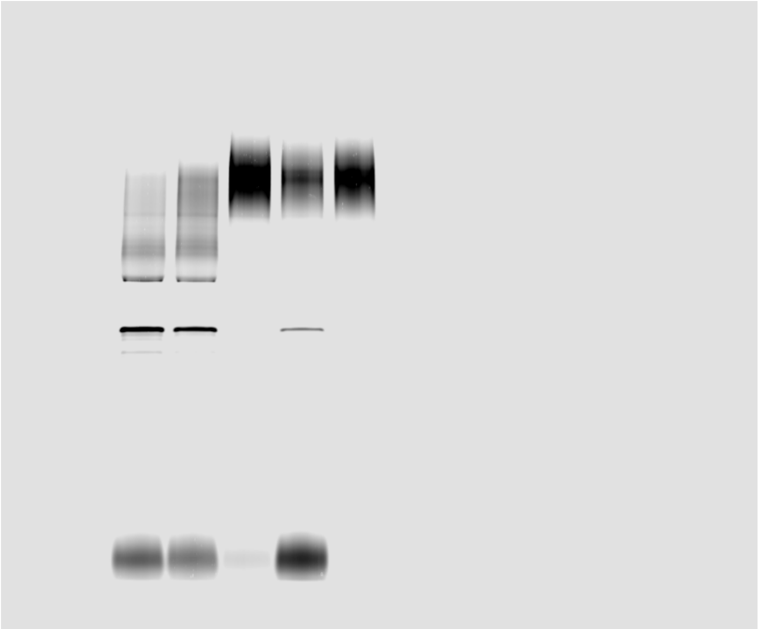

Figure 1D

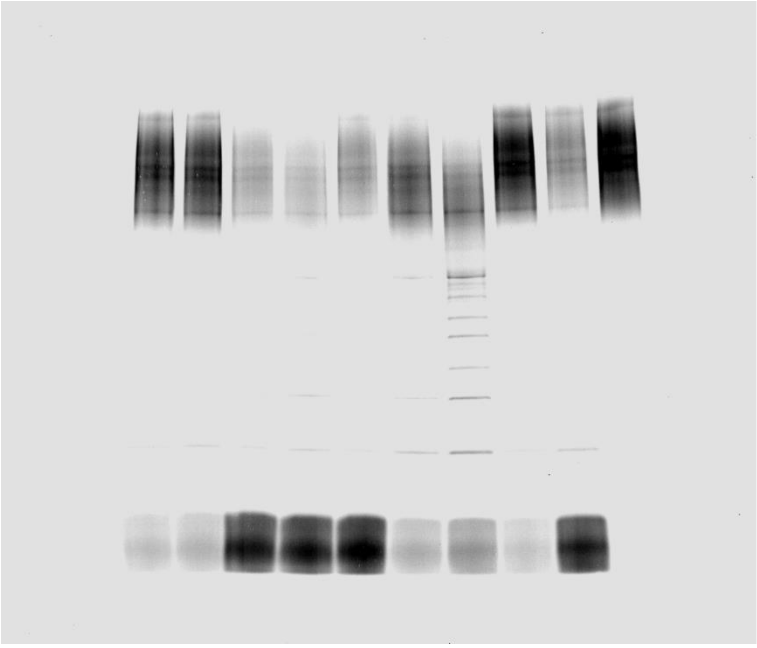

Figure 1E

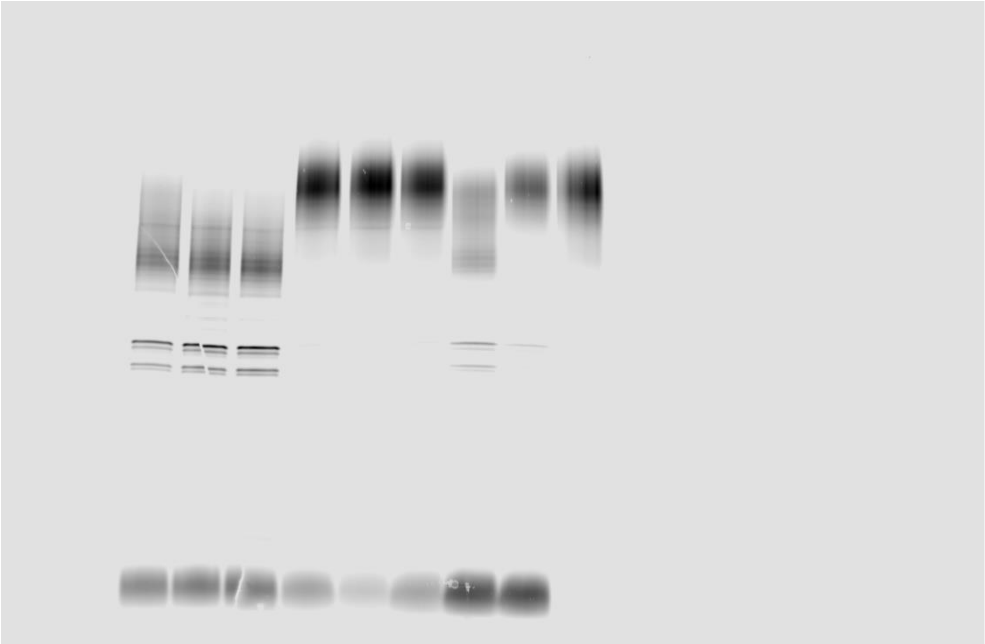

Figure 1F

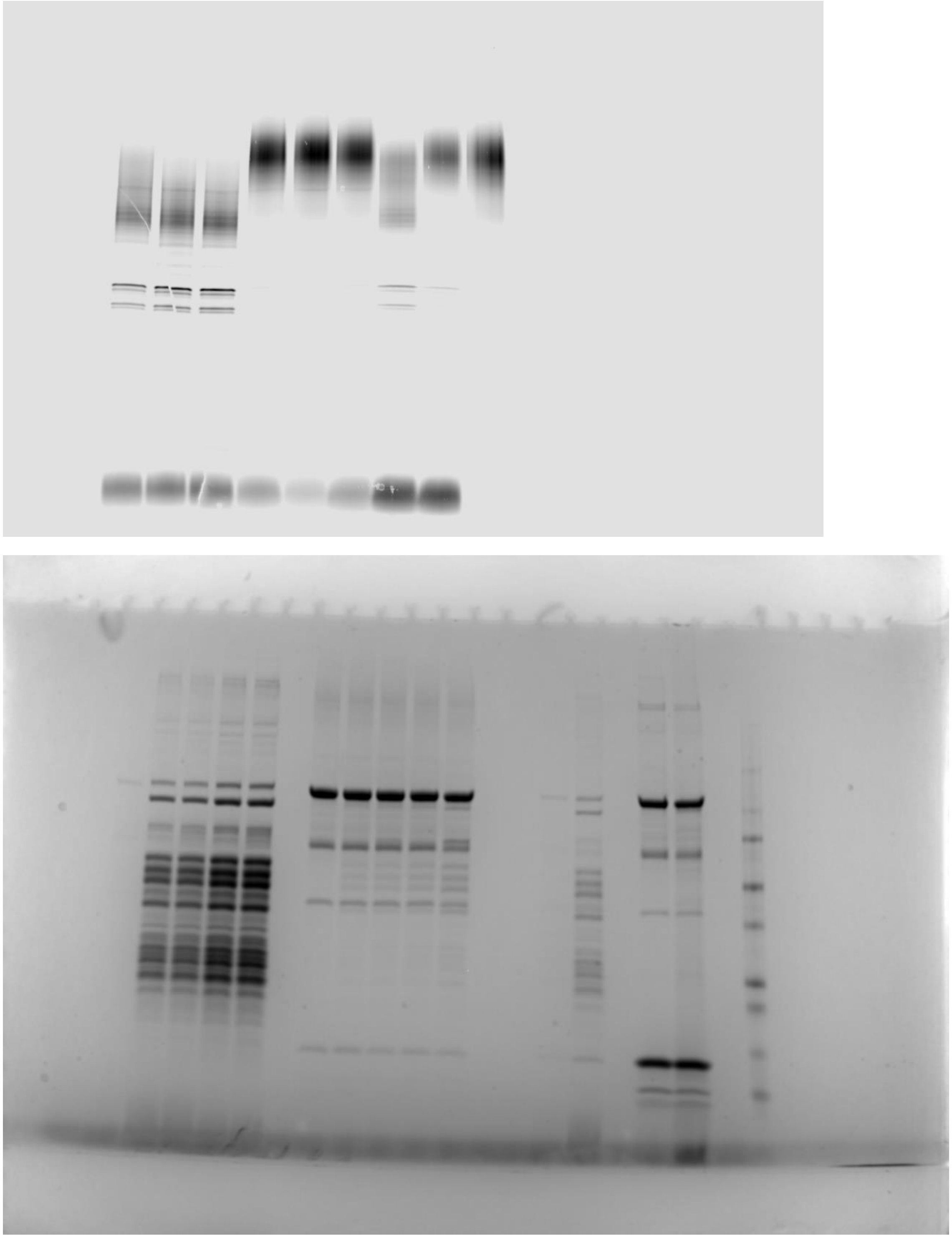

Figure 2A

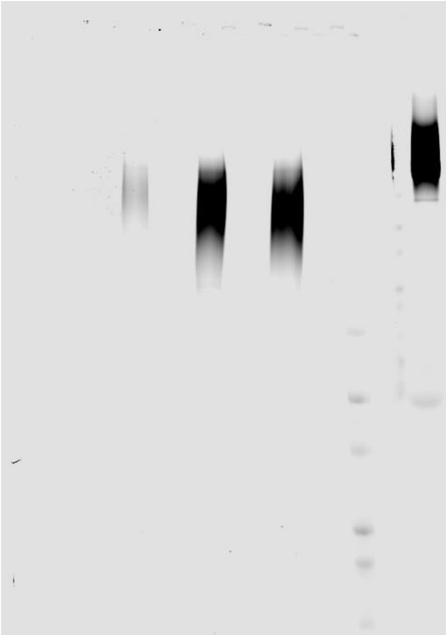

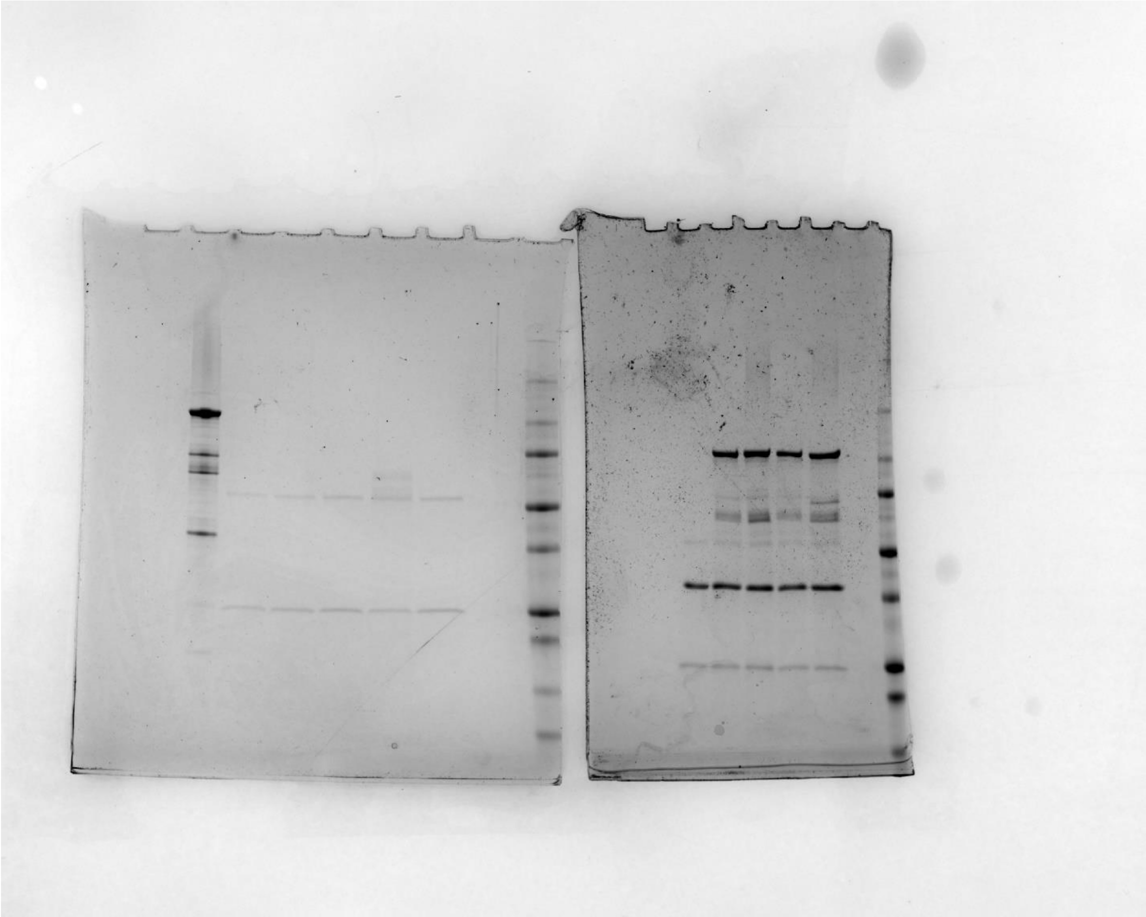

Figure 2B

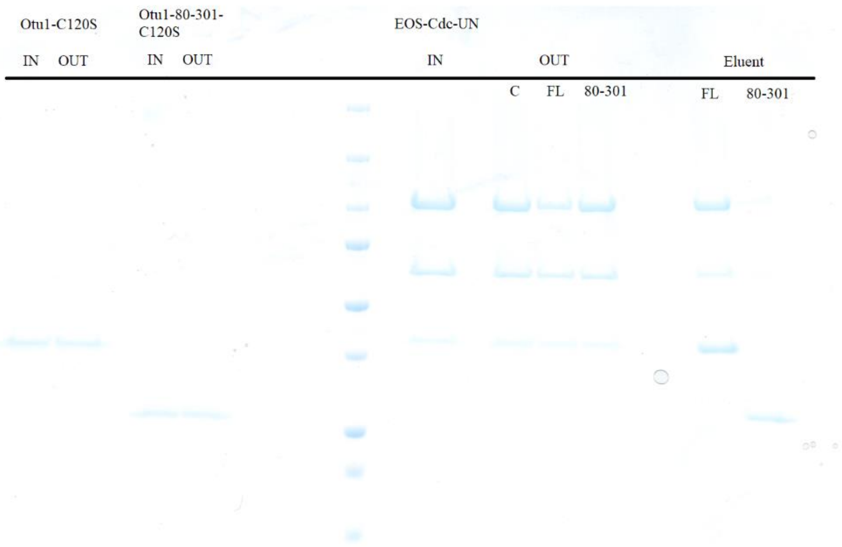

Otu1-C120S
Otu1-80-301-C120S
EOS-Cdc-UN
IN
OUT
IN
OUT
IN
OUT
Eluent
C
FL
80-301
FL
80-301

Figure 2C

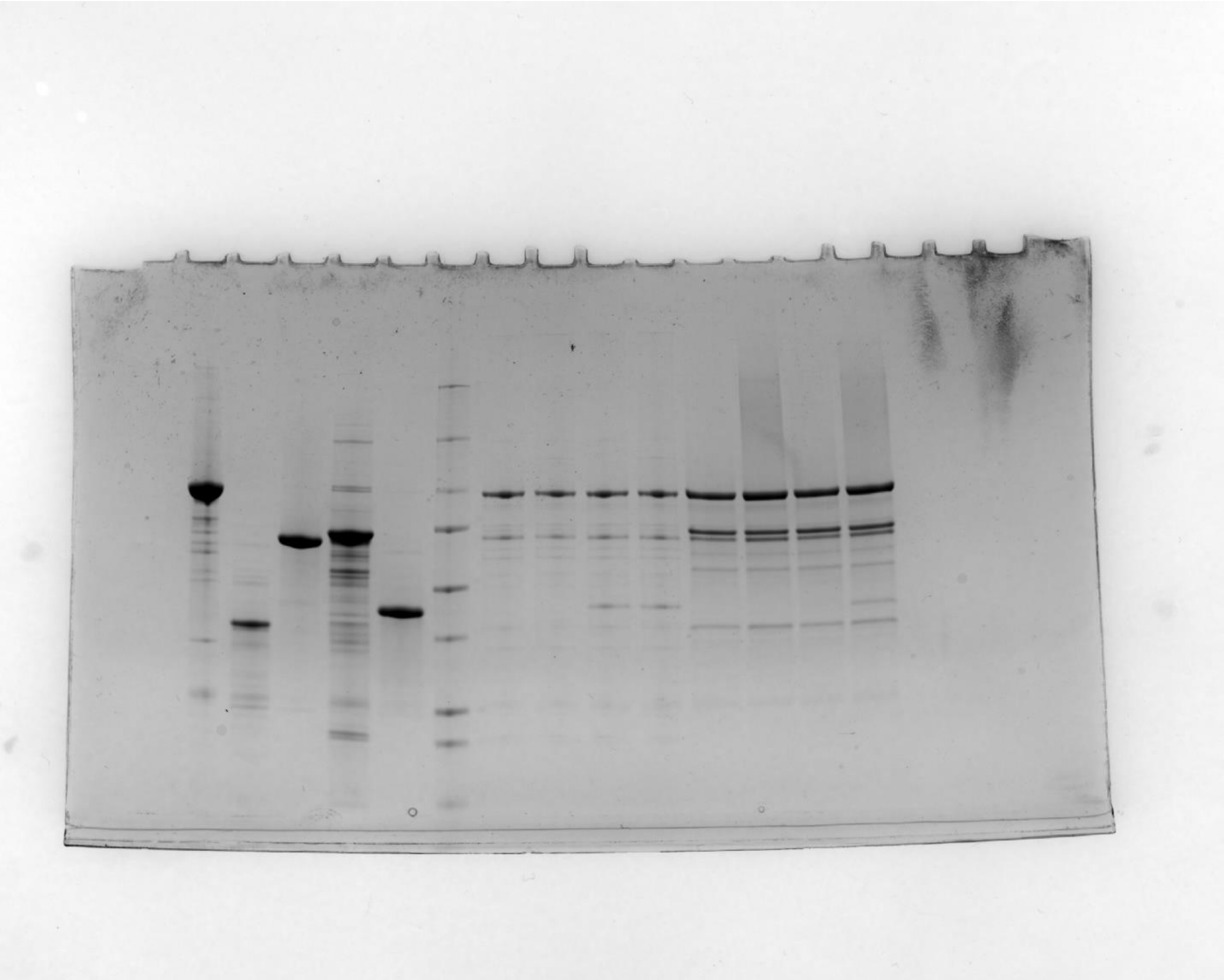
